# Fossil amber reveals springtails’ longstanding dispersal by social insects

**DOI:** 10.1101/699611

**Authors:** Robin Ninon, D’haese Cyrille, Barden Phillip

## Abstract

Dispersal is essential for terrestrial organisms living in disjunct habitats and constitutes a significant challenge for the evolution of wingless taxa. Springtails (Collembola), the sister-group of all insects (with dipluran), are reported since the Lower Devonian and thought to have originally been subterranean. The order Symphypleona is reported since the early Cretaceous with genera distributed on every continent, implying an ability to disperse over oceans although never reported in marine water contrary to other springtail orders. Despite being highly widespread, modern springtails are generally rarely reported in any kind of biotic association. Interestingly, the fossil record has provided occasional occurrences of Symphypleona attached by the antennae onto the bodies of larger arthropods. Here, we document the case of a ~16 Ma old fossil association: a winged termite and ant displaying not some, but 25 springtails attached or closely connected to the body. The collembola exhibit rare features for fossils, reflecting their courtship and phoretic behaviors. By observing the modes of attachment of springtails on different arthropods, the sex representation and ratios in springtail antennal anatomies in new and previously reported cases, we infer a likely mechanism for dispersal in Symphypleona. By revealing hidden evidence of modern springtail associations with other invertebrates such as ants and termites, new compelling assemblages of fossil springtails and the drastic increase of eusocial insects’ abundance over Cenozoic (ants/termites comprising more than the third of insects in Miocene amber), we stress that attachment with winged casts of ants and termites may have been a mechanism for the worldwide dispersal of this significant springtail lineage. Moreover, by comparing the general constraints applying to the other wingless soil-dwelling arthropods known to disperse through phoresy, we suggest biases in the collection and observation of phoretic Symphypleona related to their reflexive detachment and infer that this behavior continues today. The specific case of tree resin entrapment represents the (so far) only condition uncovering the actual dispersal mechanism of springtails - one of the oldest terrestrial arthropod lineages living today. Associations with soil-dwelling social insects over time would have been at the origin of this behavioural specialization.

## I. Background

Dispersal is essential for terrestrial arthropods living in discontinuously distributed habitats. However, acquiring dispersal features may have significant costs, such as reducing foraging efficiency for animals in confined microhabitats. In most insects, these conflicting requirements have been met through ontogenetic partitioning and the acquisition of holometaboly, allowing for soil-dwelling larvae, whereas dispersal is performed by the winged adults [1]. For wingless arthropods, the adaptive challenge of reaching new microhabitats is even greater and has led to phoretic adaptations in some arachnid groups (mites, pseudoscorpions [2, 3]).

Although considered widespread, phoresy – cases of organisms attaching onto others for the implied purpose of dispersal – is poorly understood relative to other interspecific associations [4]. As an instance of commensalism, the definition of phoresy has been revised over time to become – following the case of mites – *an association in which an organism receives an ecological or evolutionary advantage by migrating from its natal habitat while attached to an interspecific host for some portion of its lifetim*e [4–6]. The ambiguous nature of phoresy is rooted in the difficulty assessing and quantifying putative advantages over generations, whereas observable food sources (present in other commensalisms) are generally viewed as evidence of fitness improvement. Consequently, phoretic behaviors are often lent to associations between small organisms and larger mobile ones that show no evidence for other beneficial purposes. This holds especially true when trying to interpret the fossil record [7–9].

Springtails (Collembola) are entognathans, the sister-group of all other insects, and are reported since the Lower Devonian [10, 11]. The primary ecology of entognathans is thought to have been subterranean, the soil environment being suspected to have acted as an intermediate medium between water and air in the evolution of arthropods [12]. Today, springtails, which have a decisive impact on global soil microstructure and plant litter decomposition [13], are widely distributed on every continent, including Antarctica [14, 15]. As the common name suggests, their abdomen displays a jumping apparatus (the furcula) used to escape general predators. Because furcula motility is not sufficient for true dispersal, modern springtails are now only thought to disperse with soil particles as “aerial plankton” and by water for long-distance transportation [16–18]. The latter, rather unexcepted for the group, was occasionally observed and confirmed by experiments for two of the fours springtail orders (Poduromorpha and Entomobryomorpha) [17, 19, 20]. Despite their geographical and ecological ubiquity, modern springtails are very rarely reported to be engaged in biotic associations.

In addition to a few Paleozoic reports [10, 21], fossil springtails are reported from Cretaceous and Cenozoic amber deposits, in which they have occasionally been documented attached onto other invertebrates. Cases correspond to members of the order Symphypleona attached to harvestmen (Opiliones) and a false blister beetle (Oedemeridae) from the Eocene in Baltic amber [22, 23], as well as a mayfly (Leptophlebiidae) in Miocene-aged Dominican amber [24]. These springtails exhibit antennae either grasping arthropod limbs or pointing toward a potential surface of attachment, implying they were grasping onto these other arthropods while alive. This repeated fossil association has been argued to have no observed modern counterpart [22, 23].

Here, we document the case of a ~16 Ma old fossil association: a winged termite and ant with 25 springtails attached or closely connected to the body. The collembola exhibit rare features for fossils, reflecting their courtship and phoretic behaviors. By observing (1) the positions and modes of attachment of springtails on different arthropods, (2) their sex representation and (3) ratios in springtail antennal anatomies in new and previously reported cases, we infer an attachment process for dispersal in symphypleonan springtails. By revealing hidden evidence of modern springtails associated with other invertebrates such as ants and termites, new cases of fossil synclusions of Symphypleona and termites, and pointing out the drastic increase of eusocial soil insects’ ecological impact over Cenozoic (comprising more than the third of the total insect inclusions in Miocene amber), we infer that association with winged casts of ants and termites may have been a mechanism for the worldwide dispersal of these springtails. Finally, by comparing the general constraints applying to the other soil wingless arthropods known to disperse through phoresy, we evidence – aside of the specific case of tree resin entrapment – strong limits for the collect and observation of phoretic Symphypleona and make the hypothesis of their actual common occurrence in modern nature although unreported.

## II. Results

### (a) Systematic Palaeontology

Order Symphypleona Börner 1901 [25]

Suborder Sminthuridida Börner 1986 [26]

Superfamily Sminthuridoidea Börner 1906 [27]

Family Sminthurididae Börner 1906 [27]

Genus *Electrosminthuridia* gen. nov. Robin, D’Haese and Barden

#### Type species

*Electrosminthuridia helibionta* sp. nov. Robin, D’Haese and Barden

#### Diagnosis

Based on male. The genus distinguishable from other genera by combination of the following characters: antennae length about 1,8-2,5x as long as cephalic length, third and second antennomeres modified, in males, into a neat clasping organ, with a moderate angular b1 bearing at least one neat spiny setae, and a round moderate c3, fourth antennomere about 60% in male and 50-60% in females of the total antenna length, subdivided in 8 to 9 subsegments. Furcula: mucro short, very thin and tapering, ratio of mucro, dens, manubrium comparable to 1,0:1,2:0.9.

#### Derivation of name

The genus-group name is a combination of Ancient Greek, “elektron” (ἤλεκτρον) meaning ‘amber’, and *Sminthuridia* Massoud & Betsch, [28], extant genus comparable in diagnosis. The gender of the name is neutral (unstated for *Sminthuridia* Massoud & Betsch [28]).

*Electrosminthuridia helibionta* sp. nov. Robin, D’Haese and Barden

(Figs 2, 3D, SM1-2)

#### Diagnosis

As for the genus.

#### Derivation of name

The specific epithet, considered as an adjective, is a combination from the ancient Greek “helix” (ἕλιξ) describing “something twisted or spiral” and “biont”, the common internationally used suffix referring to living things. It refers to the species ability to coil its antenna around a surface to live as an epibiont of its larger host/partner.

#### Type material

Holotype, AMNH DR-NJIT001_sk (♂, Fig. 1, 2F), complete, dorsoventrally exposed, and 3 similarly exposed paratypes: AMNH DR-NJIT001_ss (♂, Fig. 1, 2E), AMNH DR-NJIT001_sj (♀, Fig. 1, 2F), and AMNH DR-NJIT001_sm (♀, Fig. 1). Five individuals (AMNH DR-NJIT001_si-m) including the holotype were extracted from main inclusion for detailed visualization.

**Figure 1.**
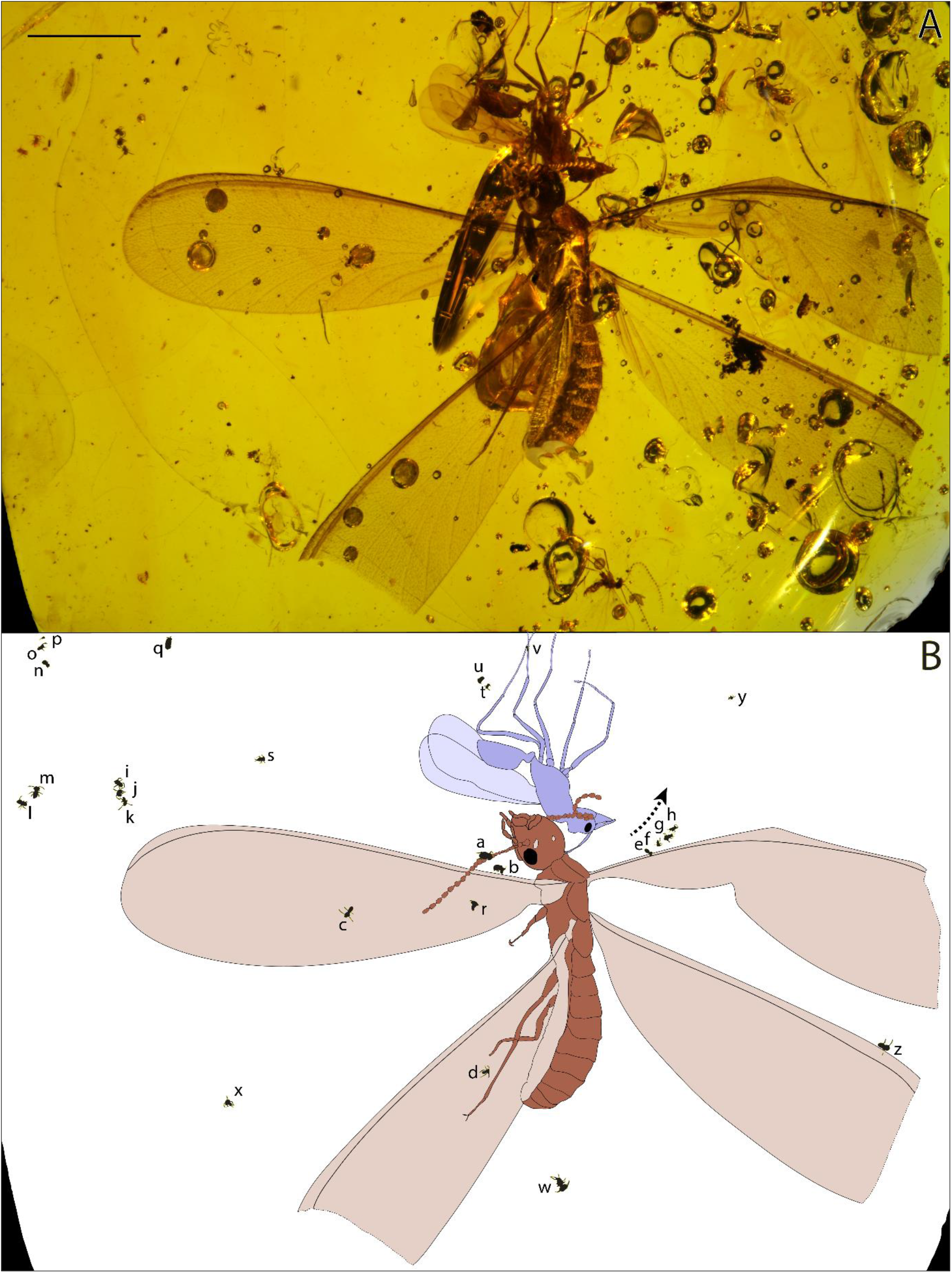
Distribution of springtails on AMNH DR-NJIT001_a-b (termite and ant) from ~16 Ma old Dominican amber. (A) Amber specimen; (B) Illustration showing the location of springtails on social insects (AMNH DR-NJIT001_sa-z). Arrow = inflow of the tree resin before consolidation. Scale bar = 0.5 cm. Photograph and interpretative drawing, N. Robin & P. Barden.

**Figure 2.**
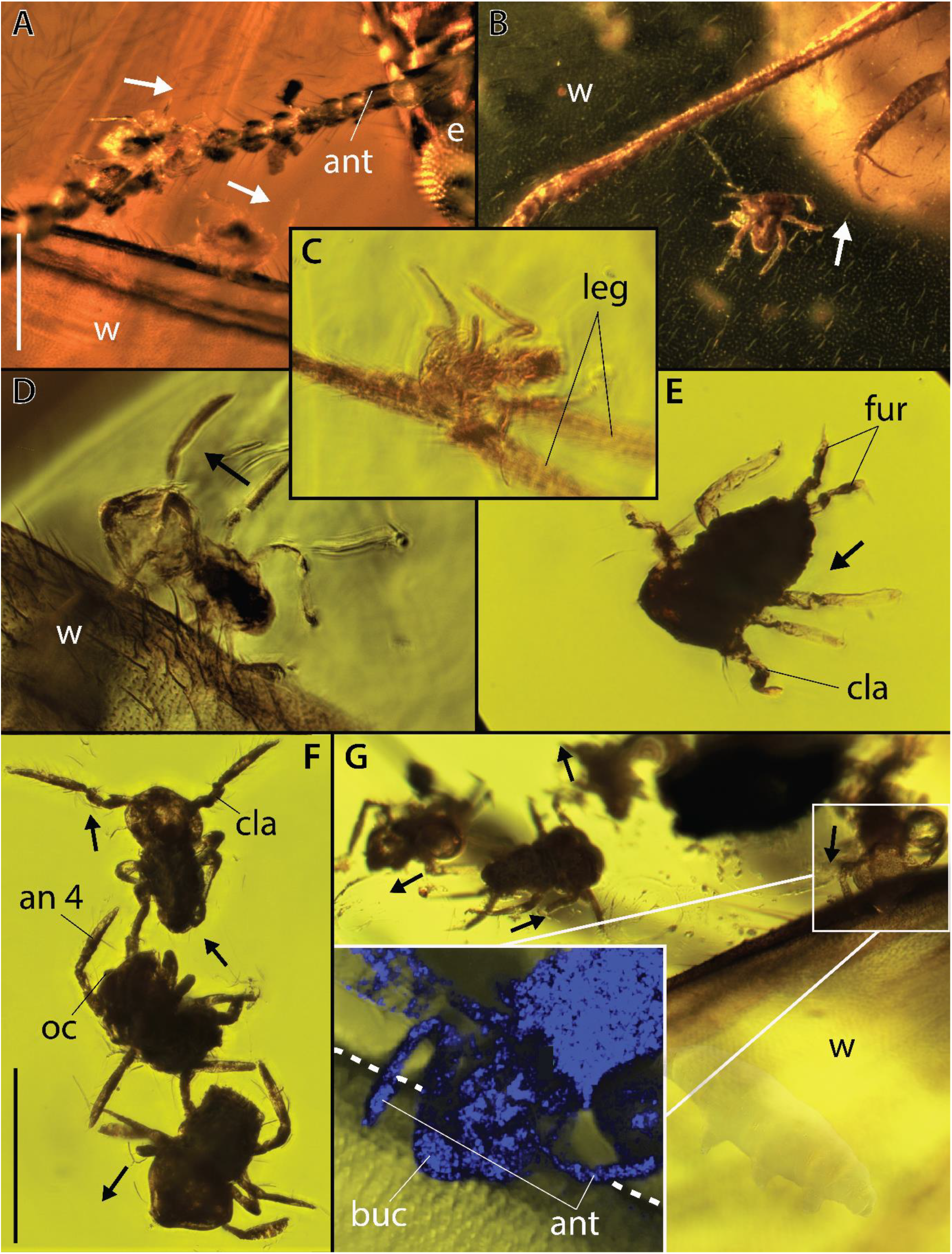
Springtails attached onto a termite and an ant (AMNHDR-0218) from ~16 Ma old Dominican amber. (A) AMNH DR-NJIT001_sa-b respectively found grasping onto the termite (AMNH DR-NJIT001_a) left antenna and left anterior wing costal margin. (B) AMNH DR-NJIT001_sd onto the termite left posterior wing. (C) AMNH DR-NJIT001_sv attached to the left second posterior leg of the ant (AMNH DR-NJIT001_b). (D) AMNH DR-NJIT001_sz found onto the termite right posterior wing costal margin. (E) AMNH DR-NJIT001_ss showing furcula. (F) AMNH DR-NJIT001_si-k displaying respectively clasping organ (AMNH DR-NJIT001_sk, male), segmented antennomere 4 and ocells (AMNH DR-NJIT001_sj). (G) AMNH DR-NJIT001_se-h found onto/close to the right anterior wing costal margin, with close-up on AMNH DR-NJIT001_se attachment to the margin using antennae and buccal cone. an = antennomere 4, ant = antennae, buc = buccal cone, e = eye, fur = furcula, oc = ocells, cla = clasping organ, leg = leg, w = wing. Arrows indicate the anterior of springtail individuals, Dashlines outlining margin costa. A-B, stereoscope images; C-G compound microscope and confocal reflectance images. Scale bars = 0.5 mm (A-B); (C-G). Photographs, N. Robin.

#### Type locality and age

Miocene amber of the Dominican Republic, La Cumbre.

#### Description

See Supplementary Material.

### (b) Fossil association

The ant/termite-springtail associations are preserved in a rectangular piece of highly transparent amber trimmed to 1.7 × 2.0 cm, also containing a male cecidomyid midge and a chalcidoid wasp. The colonized termite (AMNH DR-NJIT001_a; Fig. 1) has been identified as a *Coptotermes hirstutus†* Krishna & Grimaldi [29] (Rhinothermitidae). The ant corresponds to a dolichoderine male (AMNH DR-NJIT001_b; Fig. 1). The ant and the termite are complete and inversely oriented, both corresponding to alate reproductives. Among included springtails, 25 correspond to the sminthuridid *Electrosminthuridia helibionta* sp. nov. and one to an Isotomida (Entomobryomorpha; AMNH DR-NJIT001_sq, Fig. 1B).

One sminthuridid is found attached onto an ant leg, and seven on various regions of the termite body. The 18 other springtails are variously distributed around both of these large insects revealing key anatomical details. AMNH DR-NJIT001_sa (Fig. 1; 2A) is found on the termite left antenna with its body, head and, antenna clearly pointing toward the tenth flagellomere. Three individuals are located on the termite left hindwing (AMNH DR-NJIT001_sc, r) and forewing (AMNH DR-NJIT001_sd) membranes (Fig. 1, 2B), whereas three others are clearly restricted to the anterior margin of the termite wings. Two of them (AMNH DR-NJIT001_sb, z; Fig. 1; 2A, D) are laterally orientated onto the sclerotized anterior margin of the left hindwing (close to the wing scale) and right forewing (distally), their right and left antennae respectively overlaying this margin. This lateral posture is observed in the attachment of the single specimen found clasping onto the ant, with the left antenna overlaying the first tarsi of the ant hind leg (AMNH DR-NJIT001_sv; Fig. 1; 2C). Individual AMNH DR-NJIT001_se (Fig. 1; 2G), located onto the termite proximal right hindwing margin, shows a position even more intricately attached to the host body. Indeed, their fourth antennomere is distinctly enveloping the margin and the springtail buccal cone applied onto that surface, demonstrating a specific attachment process. Slightly distally on that margin, three other individuals are preserved in a distinct successive row, (AMNH DR-NJIT001_sf-h; Fig. 1; 2G) illustrating a linear series of detached individuals, probably trapped in the resin inflow before hardening. More generally, 14 other sminthuridids are floating in the resin close to the termite (AMNH DR-NJIT001_si-p, s, w-y; Fig. 1; 2E) and ant (AMNH DR-NJIT001_st-u; Fig. 1) some in clusters up to three individuals similarly preserved in a straight line (Fig. 2F). One specimen clearly exhibits the first undisputable clasping organ in the fossil record (Fig. 2F, 3D, SM1-2), a synapomorphy of Sminthurididae and major innovative feature allowing their specific mating behavior. This organ, specific to males, is involved in a “dancing” courtship behavior of sminthuridids, the male dragging the female onto the spot of its sperm drop by grasping its antennae [30, 31].

A total amount of 51 amber pieces from the same La Cumbre amber containing termite inclusions were screened for the presence of springtails. This piece was the only one displaying springtails, corresponding to less than 2% of our observed sampling.

## III. Discussion

### (a) Hitchhiking for dispersal

Remarkably, despite tens of thousands of known insect fossil inclusions, our report corresponds to about 6% of all the previous occurrences of springtails ever reported in Dominican amber and consist of two taxa out of twelve described in total [32–34]. As this association is unique among many comparable specimens of termites from La Cumbre, it cannot represent the general abundance of these springtails in termites’ environments but must correspond to more specific biotic explanation.

Other symphypleonan fossil associations are known from amber, beginning with two cases of Eocene harvestmen legs supporting up to five individuals arranged in a row and clasped by the antennae (Cholewinsky pers. comm. in [22]; Fig. 4). While the springtails were differently positioned (either facing the leg or facing away from it), they all show antennae distinctly bent toward this appendage, suggesting antennae were initially secured to it. The resulting position was interpreted as their immediate detachment following resin entrapment [22]. Associations with winged insects are described from attachments on the forewing base of a mayfly (Miocene [24]) and the leg of an false blaster beetle (Eocene [23]; Fig. 4C). In the case of the beetle association, antennae were implicated, as were the mouthparts which may have grasped onto the leg surface (or perhaps tibial setae, [23]). Thus, all previous cases of preserved attachment of springtails (excluding that of the mayfly association) were described from smooth cylindrical legs. The associations here reveal springtails with a positioning attachment preserved on novel structures consisting of insect antennae and wing margins. In those cases, we observe that the attachment is also achieved by rolling up of the fourth antennomere around termite antennae/sclerotized wing margins, implicating this structure in the general phoretic ability of the springtails. Possible variations in grasping mechanisms have been briefly addressed, stressing the attachments of both a long-antennomere species (*Sminthurus longicornis†*) to a 25 µm-thin leg, and that of a medium-antennomere species (*S.* sp.) to a 100 µm-thick leg [22, 23]. The herein case confirms these possible variations revealing the attachment of a much smaller species with quite short antennomeres grasped onto 20-40 µm thick structures. We stress that this behavior must have been even more dependent on antenna/body lengths. Indeed, in Poduromorpha and even more in Neelipleona morphotypes the antenna/body length ratio is less than 1:3 (Fig. 3). And if many Entomobryomorpha possess longer antennae – reaching a third to half of their total length – their limited antennal subdivision would preclude any antennal attachment that could support their total body weight (see [35], Fig. 3). Attachment efficiency may be restricted to symphypleonan morphotypes that correspond to a short bulbous body with (1) antennae reaching a third to half of the body length and a (2) fourth prehensile antennomere with subdivisions consisting at least 1/3 of the antenna length (Fig. 3). As an example, in *E. helibionta* sp. nov. the fourth antennomere is divided in eight sub-segments representing in total 60% of the total antennal length. Variations in length and subdivision of fourth antennomere among Symphypleona might have driven specificities in hosts selections/attachments, their increase conferring probably more generalist behaviors. Our observations finally confirm the involvement of mouthparts as a secondary important element to the primary antennal attachment, grasping surfaces or acting as abutment.

**Figure 3.**
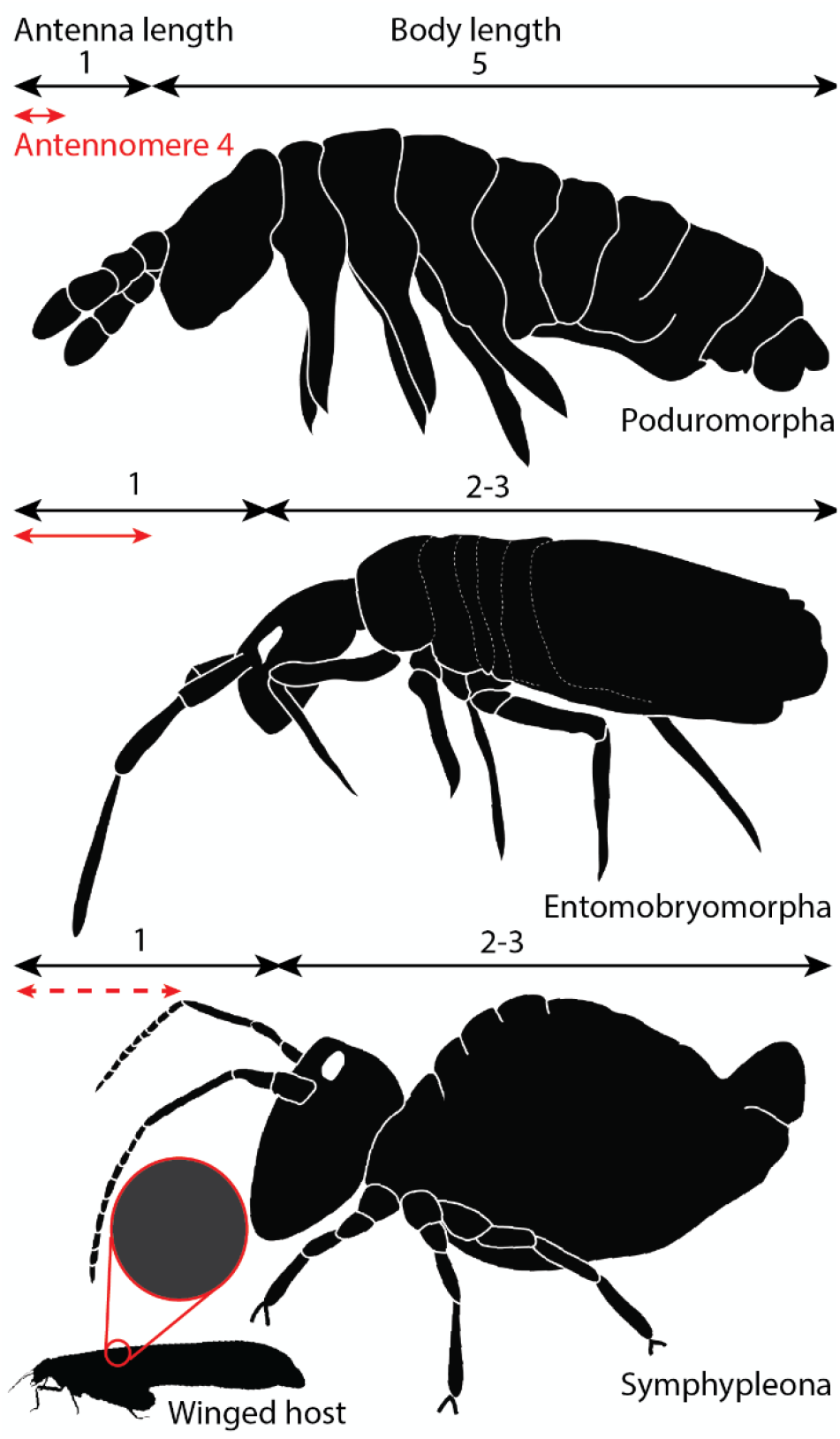
Body proportions in Collembola orders and suitability for phoretic grasping. Arrows represent anatomical lengths; figures correspond to relative anatomic ratios. Grey structure represents cylindrical support for antennal grasping on host. Dashed-line lengths represent sub-segmentation of the antenna. Neelipleona possess highly reduced antennae, they are not represented in this comparison. Drawing, N. Robin

**Figure 4.**
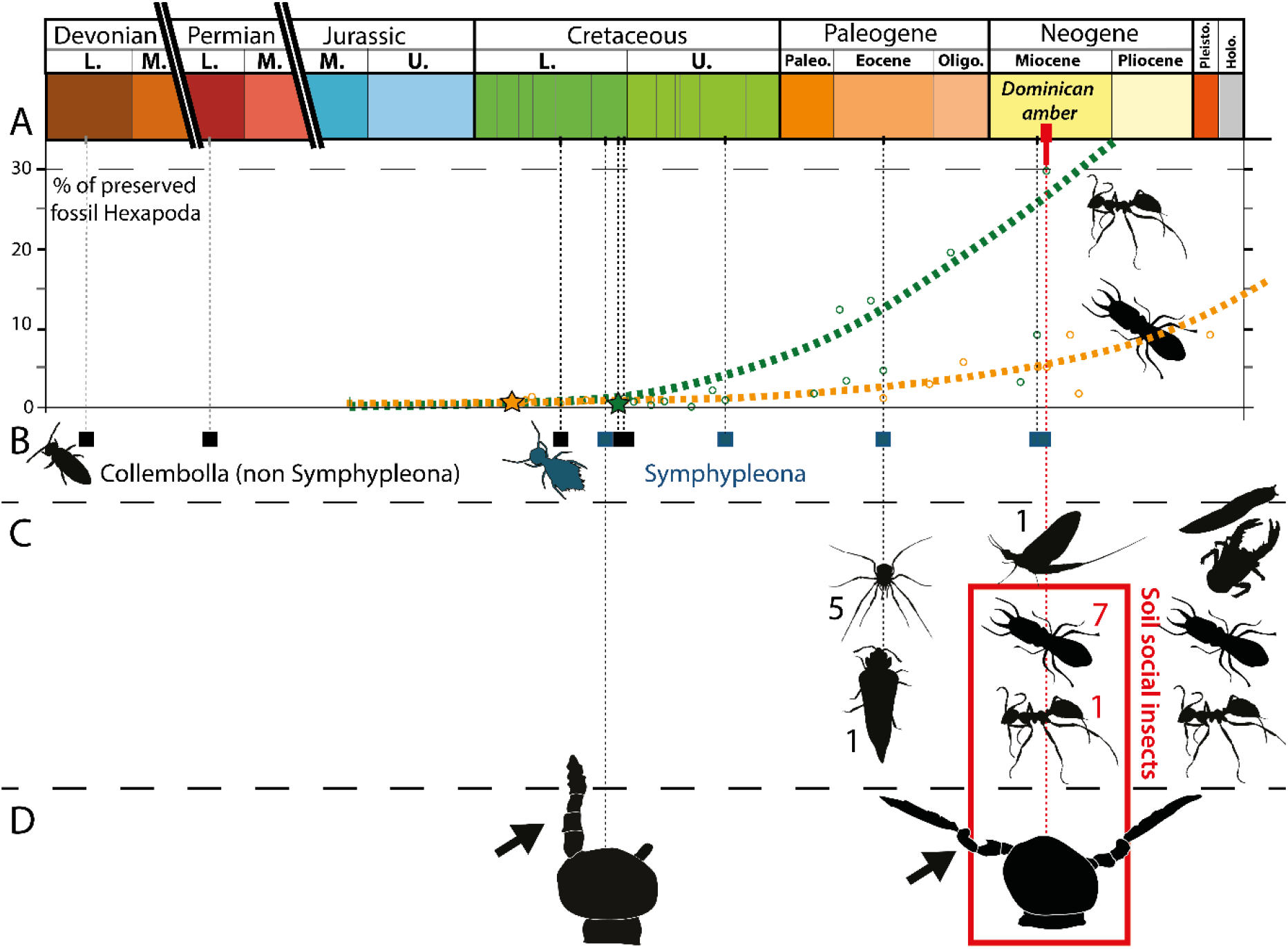
Abundance of ants and termites, collembola body fossils, phoretic behaviors and clasping organs over time. (A) Evolution of the percentage Isoptera [orange] and Formicidae [green] among other Hexapoda found from fossil amber and compression-preservation deposits. Based on abundances provided in Engel *et al.* [91] for Isoptera and Grimaldi & Agosti [92], LaPolla *et al.* [93] for Formicidae. (B) Body fossils of Collembola over time (C) Associations of Collembola onto other organisms over time, including modern cases with slugs, decapods, ants, and termites. Figures represent the amount of attached springtail individuals. (D) Clasping organs described from fossil ambers. Stars = first fossil occurrences; squares = body fossils; frame = case described here.

Antennae appear to be key for phoretic attachment, however they are subject to significant sexual dimorphism. Unlike other Symphypleona, the second and third antennomeres of male Sminthuridoidea are modified into in a clasping organ; this structure is preserved in *E. helibionta* nov. sp. However, among 25 individuals, only three males (AMNH DR-NJIT001_ss, si; Fig. 2E-F) could be identified in the inclusion. Could this disparity in sex representation have a biological underpinning? Phoretic behaviors restricted to females have been reported in other phoretic wingless arthropods. Aggregations of female mites and pseudoscorpions, sometime identified as gravids, were reported in extant fauna competing for various animal hitchhiking (harvestmen, beetle, diptera, frog) [36–39]. From laboratory experiments, Zeh & Zeh [1] even demonstrated a female bias in phoresy by pseudoscorpions that increases over time, and with mated females exhibiting a slightly higher rate of phoresy than unmated ones. Given pseudoscorpions ecological proximity and ametabolian life cycle, this – rather unexplained bias – could relate to that of phoresy among springtails observed here. The amber piece contains four distinct flying insects, some caught with wings still open, which is suggestive of an arboreal capture in the tree resin for which all trapped sminthuridids would have detached from the ant/termite association. However, the presence of one Isotomida (Entomobryomorpha) in the inclusion, could advocate for a trapping in amber close to the soil, implying the possible trapping of a few non-associated sminthuridids. Given this taphonomic limit, we cannot definitively conclude on a discrepancy in sex representation in this association. Besides, we note that comparable disparity in sex representation have been suggested for the modern symphypleonan *Sminthurides* in general sampling context [40].

While migration strategies in springtails are today poorly understood, they have been observed to occur through four vectors: windborne, pedestrian, rafting, and wind propelled on water surface [18]. Because Collembola are highly susceptible to desiccation, it is highly unlikely they are capable of moving over oceanic distances through aerial movements, favouring recently accepted water dispersal option confirmed by experiments [17, 19]. However, the so far evidence of waterborne dispersal only applies to Poduromorpha and Entomobryomorpha; as well as for large “swarm” pedestrian migrations [17, 19, 20, 41, 42]. Symphypleona have so far only been caught in air (up to 3350 m, [18, 43]) and, as strictly terrestrial and freshwater inhabitants, there is no evidence that they could maintain themselves in association with brackish water [44]. However, Symphypleona, which extends into Spanish amber dated to the Lower Cretaceous [45], display many recently diverged clades (*e.g.* genera *Sminthurinus* and *Sminthurides*) on every continent. This present-day distribution suggests the existence of additional mechanisms for dispersal. The type of phoresy evidenced herein from the fossil record may have facilitated this intensive dispersal.

### (b) Springtails and phoresy

Previous fossil reports all noted the absence of modern phoresy in springtails, highlighting a questionable discrepancy (lack of modern records vs extinct behavior, [22, 23]). Given the introduced biases affecting the identification of phoresy among commensals, we disagree with that first appraisal and uncover some, so far, hidden cases.

In fact, springtails have been reported to have associations with a diverse assortment of invertebrates, although nature of the associations has not been determined. A frequently documented case corresponds to the repeated observation of termitophile and myrmecophile inquiline springtails. Different genera of cyphoderid springtails have been observed clinging the head and back of soldiers and queen termites within nests ([46–48], Fig. 4C). In those cases, specialized sucking mouthparts, their location on soldiers’ heads and their posture – drooping toward the termite labrum – suggest that springtails obtain small meals from trophallactic soldier-worker food transfers. Cyphoderids have also been reported attached to reproductive alate ants (both females and males, [49]). Food-supply commensalism has been suggested from the observations of springtails (both Entomobryomorpha and Symphypleona) on Palearctic slugs, feeding on their mucus ([50], Fig. 4C). But springtails are also curiously reported as commensals of the shells of hermit crabs of tropical land environments ([51–57], Fig. 4C). These associations were reported from about a hundred of specimens from Mexico, New-Guinea, the Caribbean (Guadeloupe, Dominican Republic, Saint Croix). In 2005, we observed 80 individuals caught from 7 hermit-crabs and sands from different Martinican beaches (Anse Trabaud, NE of Les Salines, Grand Macabou, Presqu’ile de la Caravelle) evidencing the relative commonness of these associations within the pantropical range of these terrestrial crustaceans (Coenobitidae). The biological purpose of these associations has been addressed as a possible trophic inquilinsm [57] linked to consumption of host food remains or feces; but the actual springtails’ (Coenolatidae) location inside shells or their feeding activity was never observed. Alternatively, these associations characterize specimens collected from beach environments into which the backshore microhabitats of springtails are discontinuously distributed, requiring alternative dispersal options. Associated individuals consist mostly of females and juveniles whereas males represent about 10% of documented samples; which could be linked to an unequal sex distribution in species or a difference in sex involvement when associations relate to transport. These reports of poorly understood commensalisms with other invertebrates argue for the existence of actual phoretic-like behaviors in modern Collembola.

No modern Symphypleona is known to perform attachment to cylindrical structures by use of antennae. It has been suggested that this behavior could have been restricted to extinct lineages of Symphypleona having especially elongate flexible antennae [22]. From the variability of antennae exhibited here, we exclude this possibility. In addition, the herein fossil association reveals the nature of Symphypleona reactions to disturbances. The fossil inclusion preserves an alate termite that could not begin to fold its wings when trapped, implying an almost immediate capture in the resin. In that meantime, most springtails managed to get completely or partially detached of their host revealing the reactive mobility of the antennae, perhaps comparable to that of the furcula. A reflexive detachment may explain the apparent absence of modern phoretic Symphypleona based on modes of collection in the field. Insects and arachnids are collected most frequently by direct immersion in ethanol before being prepared for collections. It is very likely that given their superficial attachment and quick detachment system, phoretic springtails may detach from hosts in ethanol or even before immersion, effectively removing link between host and commensal (see Fig. 5a for comparison with other wingless phoretic arthropods). The tiny proportions of the fossil phoronts relative to all type of soil arthropod host (including ants and termites) size would have most of the time impeded the finding of modern representatives even when initially collected and kept in alcohol with that host (Fig. 5b). In this context, the immediate embedding of springtails onto insects by tree resin would characterize a unique situation enabling the preservation of these associations, making the fossils of high significance to document and study them.

**Figure 5.**
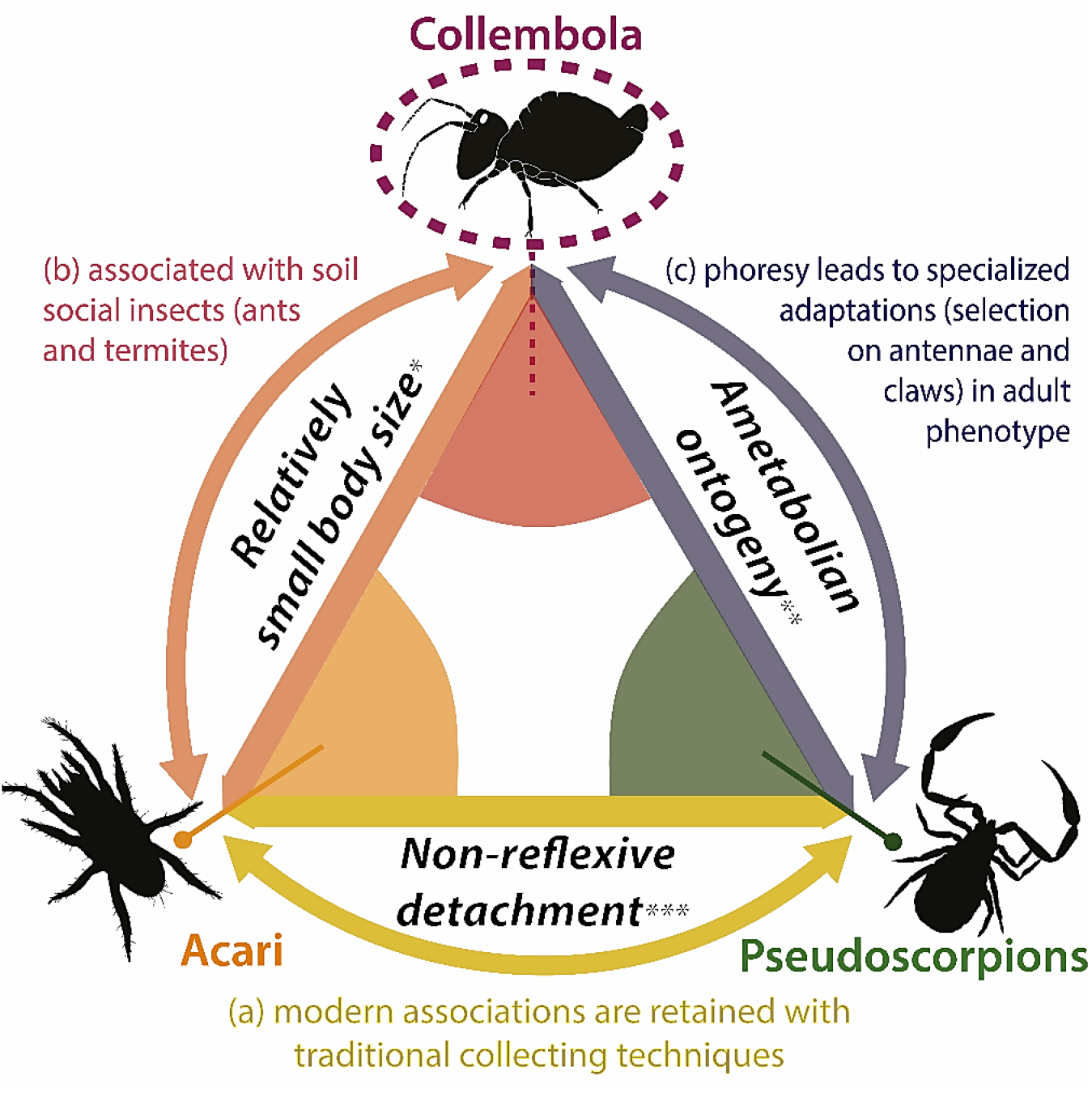
Morphological, ontogenetic and behavioral constraints for phoretic strategies among apterous soil-dwelling arthropods. Three criteria shape the phoretic interactions of apterous arthropods with larger hosts, introducing the new case of collembola (Symphypleona). Cases of collembolan (symphypleonan) phoresy correspond with a specific set of conditions that are not met by other phoretic soil-dwellers. Criteria support the heterogeneous nature of reports of these different associations in modern instances, with the so far totally unreported case of modern Symphypleona associations. * much smaller relative to all types of other soil arthropods; ** no specific phoretic lifestage during ontogeny; *** of claws or mouthparts.

### (c) Evolving with social insects

Three of the four orders of modern springtails are reported from termite and ant nests [48, 58–60]. This includes cyphoderids primarily described as nests inquilines as well as direct commensals of termites and ants [46–49, 60]. Thus, despite limits in collecting modern phoretic springtails, other reported cases suggest significant relationships with social insects. Moreover, we report paleoecological insight that further bolsters links between springtails and social insects. Based on an initial screening of more than 1300 inclusions within rough Cambay amber, from the Lower Eocene (54 Ma) of western India, we report only two Symphypleona. Those two individuals, representing different genera, are both located a few millimeters from a single alate termite.

Today, ants and termites together comprise a significant component of many terrestrial ecosystems. The biomass of termites is estimated to be approximately equal to that of humans [61] and in tropical localities, termites and ants together may outweigh all vertebrates and all other insects combined [62]. It is possible to trace the “rise” of soil-dwelling social insects from their first appearance in the fossil record to their remarkable ecological impact today. The termite and ant fossil records extend to the Lower Cretaceous of Russia (Berriasian) and Charentese-Burmese ambers (Albian, Fig. 4A), respectively. In the Cretaceous, ants and termites never comprise more than 2% of all fossil hexapods by locality [63, 64]. This changes markedly by the Cenozoic. In Dominican amber, termites make up to 6% of inclusions while ants represent ~30% of all hexapods; social insects represent more than 1/3 of the total entomofauna at that time (Fig. 4A). As the vast majority of soil insects, termites and ants stand as an immediate model for widespread biological interactions in springtails.

Through various modern ecologies (soil, leaf litter, canopy), ant lineages are primarily soil or surface dwelling [65]. For Blattodea – *ie*. cockroaches and termites – modern groups live in the superficial soil including litter and in wood logs, corresponding to their ancestral ecology [66, 67]. The host termite identified here, *Coptotermes* is also distinctly subterranean [66–68]. Advanced levels of sociality are documented in fossil stem-ants as early as the mid-Cretaceous [50,51] and must have applied to the roaches-termites lineage since the early Mesozoic [43, 44], implying that the organisation in nests was present early in the emergence of these two taxa.

Consequently, various springtail families have been confronted with the fast-increasing prevalence of eusocial insects from the mid-Cretaceous onwards, eventually leading to advantages of living close or inside their nests, as reported for most orders of springtails ([48, 58–60], Fig. 4B). Termito/myrmecophile ancient behaviors have previously been suspected for other Dominican amber springtails (cyphoderines, [32]).

Living in close proximity to termites and ants, as they increased in ecological impact, would provide significant benefits for springtails. An initial termito/myrmecophile behavior would then act as a catalyst, providing opportunity for the evolution of more specialized associations. This type of specialization to eusocial environment is actually also observed in other groups of early diverging hexapods like bristletails (Thysanura), which are reported as myrmecophiles [73] but also as direct commensals climbing on large ant larvae at colony migration [74]. Thus, beside termito/myrmecophile behaviors, associations with eusocial insects would have appeared at least twice in the evolution of springtails: in cyphoderines (Entomobryomorpha) for apparent feeding and in Symphypleona for (primary or secondary) dispersal. From both fossil and modern examples, we infer that social insects would have acted as Symphypleona dispersal agents. The phoresy would have been enabled through hitchhiking on alate termite/ants adults, at the time they reach the leaf litter to begin their nuptial flights. Comparable to many symphypleonan genera, *Coptotermes* has distributed onto every continent with diversification since just the Miocene [75], implying its strong ability for overseas dispersal. As for dispersal, modern subterranean termites showed real flight performances reaching a 900 m distance in a single take-off and crossing in the same time very large water mass (e.g. the Mississippi river, [76]). The grasping ability of Symphypleona would have been enhanced by the elongation/segmentation of the fourth antennomere. This derived feature may have allowed the hitchhiking of other soil surface inhabitants (toward more or less efficient dispersal), as observed in other fossil examples (Fig. 4C).

From both fossil and modern cases, soil-dwelling social insects show an obvious trend for biotic associations with smaller apterous arthropods. Indeed, modern mites (both parasitiforms and acariforms) are known for living in ant nests, with phoretic nymphs found on extant ant alates, as well as army ant workers [43, 59, 60]. These nymphs, called hypopi, correspond to a non-feeding stage of mite ontogeny, specialized for phoresy. These associations have been reported twice as early as 44 Ma (Mesostigmata; 56,57). If less described, termitophilous mites are also abundant [77, 81, 82], with not less than eight (parasitiform and acariform) families detected from the study of only three subterranean termites (including *Coptotermes*) in various continents [63–67]. Limits on these associations remain the size of soil social insects, restricting phoresy to mites and other millimeter-sized organisms, such as springtails (Fig. 5b). Contrary to mites, the ontogeny of springtails displays no metamorphosis (hexapods ametaboly, [88]) and thus, no possible specific stage to increase dispersal efficiency. It is therefore relevant that phoresy in this group appears in the adult-phenotype (through attachment structures) and primarily in lineages most armed for this kind of acquisition (Symphypleona). Springtails share this criterion with pseudoscorpions in which phoretic abilities are displayed by the adult phenotype as well, namely through adapted claws (Fig. 5b). In this regard, springtails phoresy compares with the selection constraints of the attachment strategy found in pseudoscorpions, applying within the host range found in mites, but controlled by a much more reactive behavior, probably reflective of hexapod motility speed compared to that of most arachnid orders ([89], Fig. 5). Through their body proportions and antennae length, Symphypleona would have checked the requirements for specialized phoretic behaviors among other springtail morphologies (Fig. 3). Limited understanding of modern counterparts impedes consideration of whether these phoretic behaviors could have shaped the Symphypleona general morphology as well.

## IV. Conclusion

Our results detail the existence of a new type of phoretic behavior among wingless soil arthropods: the one displayed by springtails. This behavior would have been at the origin of the actual worlwide dispersal of a significant member of springtail (order Symphypleona) never reported from marine water and therefore unlikely to have followed the long-term dispersal process employed by other lineages. Our survey of hidden associations of springtails with larger organisms argue for their actual ability to set up generalist and specialized relationships with several hosts, although these are almost unstudied. Fossil and modern occurrences support a connection between the apperance of phoresy among springtails and the increase of social insects’ ecological impact over time. Their great abundance, minute size and social organisation including (1) soil castes sharing the early habitat of Symphypleona, and (2) winged casts flying towards different microhabitats at mating, would have constituted a key vector for dispersal strategy in that group – one of the oldest living terrestrial arthropod lineages. This mode of phoresy refers to selection pressures applied onto the adult phenotype as springtails undergo no metamorphosis throughout their ontogeny, as in pseudoscorpions. Their potential range of host sizes is comparable with that observed for mites. However, as hexapods, springtails movement speed (as well as amber entrapments evidences) would argue for a system of detachment so quick that traditional arthropod sampling methods would have so far missed modern occurrences. We propose that these associations remain common in extant taxa, albeit hidden until alternative sampling techniques are employed.

## V. Methods

The amber specimen hosting the association (AMNH DR-NJIT001) originates from La Cumbre, Dominican Republic and is deposited at the Division of Invertebrate Zoology at the American Museum of Natural History. The amber has been trimmed and polished with a water-fed trim saw and Buehler EcoMet™ 30 grinder to improve imaging and reveal additional detail. Lower magnification (1-2x) extended-focus photomicrographs were taken with a Nikon SMZ-18S through NIS Element D software. Higher magnification pictures were captured with a Nikon Eclipse TS2 compound microscope after immersion in glycerol. Reflectance images of springtails were captured with a Leica SP8 confocal scope (NJIT) and processed using the LAS X software. For systematics, we follow Derhaveng [90] for more inclusive ranks and Bretfeld [40] for subordinal classification. For generic assignment we build upon Massoud & Betsche [28] which includes the most complete comparative work of secondary sexual characters in modern Symphypleona.

## V. Authors contributions

NR and PB have collaboratively performed amber specimen preparation and imaging, as well as drafted the general manuscript. PB provided the report of the Cambay amber synclusion of springtails with an alate termite. Drawings and schemes were done by NR. CDH provided novel observations on hidden cases of modern springtail associations and supervised taxonomic assignments.

## VI. Acknowledgments

We are grateful to Viktor Baranov for identifying the associated gall midge. We thank Jorge Martinez and Keith Luzzi for facilitating access to specimens. Hukam Singh and David Grimaldi for access to Cambay amber specimens and proportions of arthropod inclusions. N. Robin thanks L. Legender for discussion on taxa names. This work has been achieved along a Fulbright Visiting Scholar Program (National Program) of the French-American Commission (*Tracking 100 million years of biological symbioses in fossil termites* – N. Robin).

