## Supplementary Material for "Fossil amber reveals springtails’ longstanding dispersal by social insects"

Order Symphypleona Börner 1901 [1]

Suborder Sminthuridida Börner 1986 [2]

Superfamily Sminthuridoidea Börner 1906 [3]

Family Sminthurididae Börner 1906 [3]

Genus *Electrosminthuridia* gen. nov. Robin, D’Haese and Barden

**Type species**. *Electrosminthuridia helibionta* sp. nov. Robin, D’Haese and Barden

**Diagnosis.** Based on male. The genus distinguishable from other genera by combination of the following characters: antennae length about 1,8-2,5x as long as cephalic length, third and second antennomeres modified, in males, into a neat clasping organ, with a moderate angular b1 bearing at least one neat spiny setae, and a round moderate c3, fourth antennomere about 60% in male and 50-60% in females of the total antenna length, subdivided in 8 to 9 subsegments. Furcula. mucro short, very thin and tapering, ratio of mucro, dens, manubrium comparable to 1,0:1,2:0.9.

**Derivation of name.** The genus-group name is a combination of Ancient Greek, “elektron” (ἤλεκτρον) meaning ‘amber’, and *Sminthuridia* Massoud & Betsch, [4], extant genus comparable in diagnosis. The gender of the name is neutral (unstated for *Sminthuridia* Massoud & Betsch [4]).

*Electrosminthuridia helibionta* sp. nov. Robin, D’Haese and Barden

(Figs 2, SM)

**Diagnosis.** As for the genus.

**Derivation of name.** The specific epithet, considered as an adjective, is a combination from the ancient Greek “helix” (ἕλιξ) describing “something twisted or spiral” and “biont”, the common internationally used suffix referring to living things. It refers to the species ability to coil its antenna around a surface to live as an epibiont of its larger host/partner.

**Type material**. Holotype, AMNH DR-NJIT001_sk (♂, Fig. 1, 2F), complete, dorsoventrally exposed, and 3 similarly exposed paratypes: AMNH DR-NJIT001_ss (♂, Fig. 1, 2E), AMNH DR-NJIT001_sj (♀, Fig. 1, 2F), and AMNH DR-NJIT001_sm (♀, Fig. 1). Five individuals (AMNH DR-NJIT001_si-m) including the holotype were extracted from main inclusion for detailed visualization.

**Additional examined material.** The other synincluded 21 (more or less complete) representatives of the species.

**Type locality and age.** Miocene amber of the Dominican Republic, La Cumbre.

**Description.** (from type material)

**Male**. Total length as preserved (from the tip of the head to tip of the opisthosoma): 480 µm. Head length as preserved: 210 µm, width: 200 µm, cephalic diagonal not accessible in male types, with moderate short setae; eyes not visible in male types; antennae length: 620 µm, about 2,4x as long as cephalic length, with a segments length ratio 1:1,5:1,3:4,7; third and second antennomeres modified into a neat clasping organ as figured (Figs 2, SM), with a moderate angular b1 bearing at least one neat spiny setae, fine noticeable b2, b3, b4, and a round moderate c3 (giving a cubit shape to the antennomere 3) bearing at least 2 long spiny setae and smaller ones anteriorly; trichobothria not visible, fourth antennomere, about 60% of the total antenna length, subdivided in 8 to 9 subsegments, at least three long setae (about 8 µm) on each. Legs. Coxa not visible in male types; trochanter length: 50 µm; femur length: 110 µm; tibiotarsus length: 140 and up to 155 µm on hindlegs; tibiotarsal organ not visible; unguis not visible in male types. Collophore. Not visible. Retinaculum. Not visible. Opisthosoma. Length 290 µm as preserved; thorax segmentation slightly visible; greater abdomen with two/three subsegments distinguishable posteriorly and with posterior margin quite angular, lesser abdomen about a quarter in width and round; bothriotrichia not visible. Furcula. About a third of the body total length; manubrium length: 63 µm; dens length: 86 µm, mucro short, very thin and tapering, length 68 µm, broad, width 13 µm, mucronal lamellae not visible. Ratio of mucro, dens, manubrium: 1,0:1,2:0.9.

**Female.** Total length as preserved (from the tip of the head to the tip of the opisthosoma): 530 µm. Head length as preserved: 258 µm, width: 220 µm, cephalic diagonal: 340 µm, with moderate short setae; a pair of symmetric barb-like short extensions visible on some specimen mouthparts; eyes almost never visible, with 3-4 ommatidia from a unique visible right eyepatch; antennae length: 443 µm, about 1,8x as long as cephalic length, with a segments length ratio 1:1,4:1,14:5,2; trichobothria not visible, fourth antennomere, about 50-60% of the total antenna length, subdivided in 8 subsegments, at least three long setae (about 80 µm) on each. Legs. Coxa length: about 4 µm as preserved; trochanter length: 5,6-7,2 µm on hindlegs; femur length: 67-90 µm on hindlegs; tibiotarsus length: 160-186 µm on hindlegs; tibiotarsal organ not visible; unguis elongate, length: 30-55 µm, about 0.25x length of tibiotarsus, slightly curved with thin hook very distally, unguiculus present but poorly visible, about as long as unguis. Collophore. Not visible. Retinaculum. Not visible. Opisthosoma. Length 370 µm as preserved; thorax segmentation visible; greater abdomen with two/three subsegments distinguishable posteriorly and with posterior margin quite angular, lesser abdomen about a quarter in width and round; bothriotrichia not visible; anal valvules slightly visible, without trace of subanal appendages. Furcula. Not visible on female types.

**Remarks.** Specimens display a straightforward Symphypleona overall morphology with indistinct abdominal segments and globular appearance. The modification of antennomeres II and III in a neat clasping organ in male representatives (the first undisputable fossil version) for courtship behaviour, allows their placement into the Sminthuridida Börner [2] that contains two families. The monospecific Mackenziellidae Yosii [5], displaying very different elongated bodies, can be excluded placing the species into Sminthurididae Börner 1906. Sminthurididae currently comprise 11 extant and one fossil genera. The herein described material is represented both by adult (differentiated) female and male individuals. By displaying a systematic subsegmentation of the 4^th^ antennomere in both sexes, the material distinguish from [*Stenacidia*](https://fr.wikipedia.org/w/index.php?title=Stenacidia&action=edit&redlink=1) Börner [3], *Sphaeridia* Linnianemi [6], *Sminthurides* Linnianemi [6]*,* [*Denisiella*](https://fr.wikipedia.org/w/index.php?title=Denisiella&action=edit&redlink=1) Folsom & Mills [7] (redescribed in Palacios-Vargas et al. [8]) and [*Boernerides*](https://fr.wikipedia.org/w/index.php?title=Boernerides&action=edit&redlink=1) Bretfeld [9]. It also differs from the quite original monospecific genera *Sinnamarides* Betsch & Waller [10], *Pedonides* Bretfeld [11] and *Pseudosminthurides* † Sanchez-Garcia & Engel [12] in exhibiting respectively short head and mouthparts, no clasping structures on leg II and visibly modified antennomeres II and III, in addition to a common small size for the family (<0.5mm whereas *Pseudosminthurides* exceeds that size). From observable chaetotaxic elements of the male clasping organs, the genus distinguishes from [*Debouttevillea*](https://fr.wikipedia.org/w/index.php?title=Debouttevillea&action=edit&redlink=1) Murphy [13] and [*Pygicornides*](https://fr.wikipedia.org/w/index.php?title=Pygicornides&action=edit&redlink=1) Betsch [14] by displaying respectively no modification of b and c elements (antennomeres II and III) into vesicles or lamellae. The clasping organ anatomy evokes that of [*Sminthuridia*](https://fr.wikipedia.org/w/index.php?title=Sminthuridia&action=edit&redlink=1) Massoud & Betsch [4] and [*Yosiides*](https://fr.wikipedia.org/w/index.php?title=Yosiides&action=edit&redlink=1) Massoud & Betsch [4] placing into the range of what these authors describe as poorly to moderately modified antennae, and for which they note the absence of additional secondary sexual characters. Present species distinguishes from [*Yosiides*](https://fr.wikipedia.org/w/index.php?title=Yosiides&action=edit&redlink=1) in having a neat tapering mucron. It does not sharply differs from the short diagnosis of [*Sminthuridia*](https://fr.wikipedia.org/w/index.php?title=Sminthuridia&action=edit&redlink=1). Given the fossil condition of the present material, precluding accurate visualization of a possible tibiotarsal organ, mouthparts, collophore, retinaculum and detailed chaetotaxy for trichobothria in clasping organ and setae in general, we assign the present material to the monospecific new genus *Electrosminthuridia* to emphasize its proximity to *Sminthuridia* from an incomplete anatomical point of view, as previously proceeded by Sanchez-Garcia & Engel [12] with *Pseudosminthurides* †. If antennomeres II and III of *Electrosminthuridia helibionta* sp. nov. are clearly modified in a clasping organ, we notice that the usual dimorphism observed in modern Sminthurididae seems quite reduced here, with males about the same size than females. Diagnoses were taken from Massoud & Betsch [4] comparative work on Sminthurididae clasping organs [14], complemented by the original diagnoses of subsequently described genera (*Sinnamarides, Pedonides, Pseudosminthurides*) as well as key generic features provided in Bretfeld [15] for original diagnoses relying on the sole description of the clasping organ (eg. *Sminthuridia* and *Yosiides*).

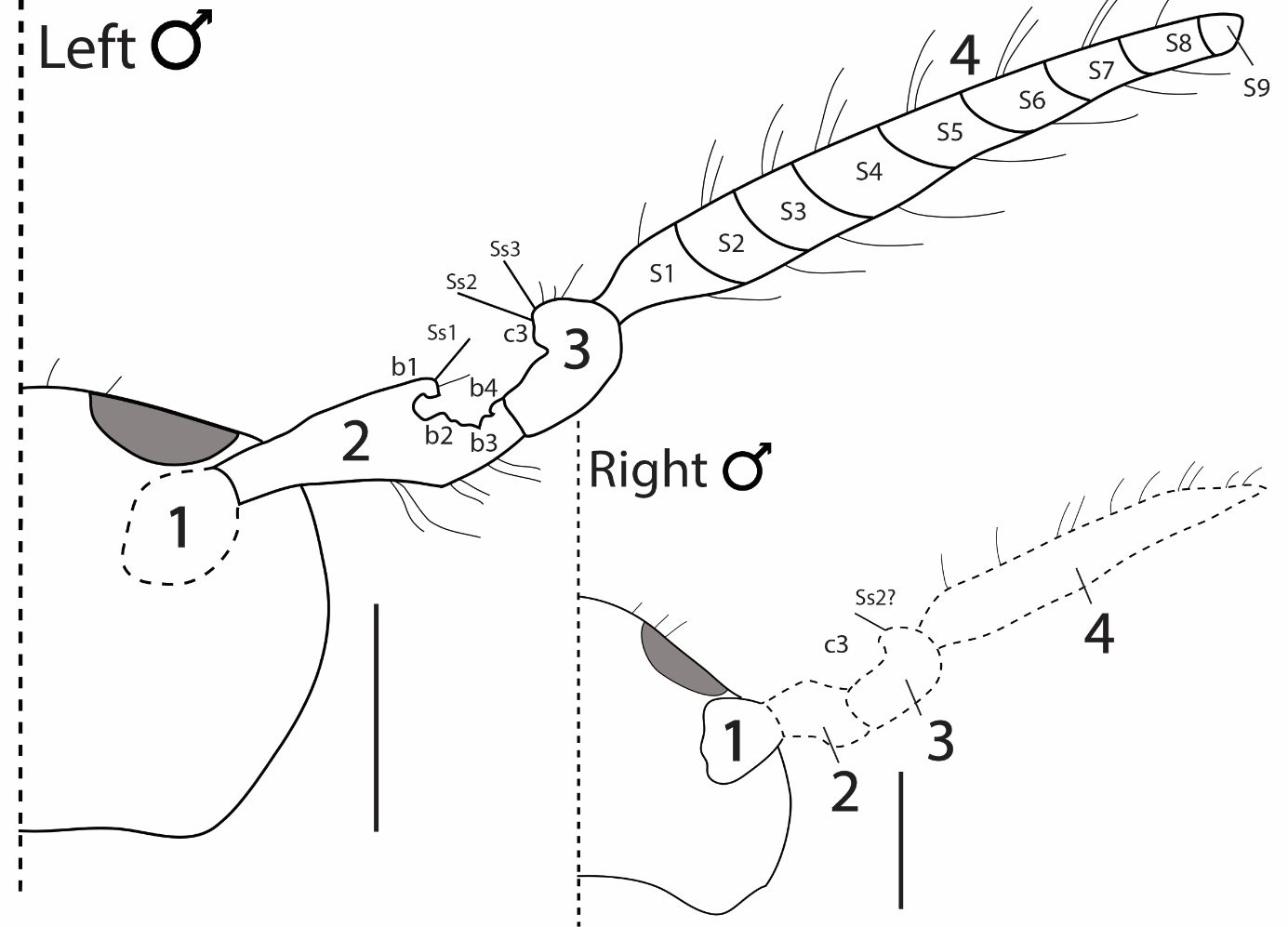

**Figure SM1.** Interpretative drawing of the male clasping organ chaetotaxy and antennal structure in *Electrosminthuridia helibionta* sp. nov. (from AMNH DR-NJIT001_sk; ♂). 1-4 = antennomere 1-4. b1-4 = specific elements of antennomere 2, c3 = specific elements of antennomere 3, S1-9 = subdivisions of antennomere 4, Ss 1-3 = spiny setae on antennomeres 2 & 3. Scale bars = 0.01 mm. Drawing, N. Robin.
